# Differential gene expression reflects larval development and survival of monarch butterflies on different milkweed hosts

**DOI:** 10.1101/2020.09.05.284489

**Authors:** Pablo M. Gonzalez-De-la-Rosa, Mariana Ramirez Loustalot-Laclette, Cei Abreu-Goodger, Therese Ann Markow

## Abstract

Second instar larvae of the monarch butterfly, *Danaus plexippus*, from a nonmigratory population in Irapuato, Mexico, were reared for twenty-four hours on three species of milkweed hosts: *Asclepias curassavica, A. linaria, and Gomphocarpus physocarpus.* We then measured larval growth and differential expression of coding genes and of microRNAs. Larval growth was similar on the two *Asclepias* species, while little growth was observed on *G. physocarpus*. The greatest differences in coding gene expression occurred in genes controlling growth and detoxification and were most extreme in comparisons between *G. physocarpu*s and the two *Asclepias*. MicroRNAs are predicted to be involved as regulators of many of these processes, in particular miR-278, differentially expressed here, could be an important regulator of growth through Hippo signaling. The implications for survival of the monarch, especially in the context of environmental factors altering the availability of their favored milkweed species, are discussed.

## 1. INTRODUCTION

Monarch butterflies, *Danaus plexippus,* are famous for their annual long-distance migrations from overwintering sites in central Mexico to as far north as Canada. The migrations involve three to five generations before the return in the fall to overwinter in oyamel or “sacred fir” (*Abies religiosa*), trees in the Monarch Butterfly Biosphere Reserves (MBBR) in the states of Mexico and Michoacan. In addition to this large Eastern population is another, the Western Population, in California, that undergoes a shorter migration, as well as a number of non-migratory populations in places such as Florida, Hawaii and Guanajuato, Mexico (Pierce et al., 2014; Zhan et al., 2014). Monarch populations are on the decline (Vidal & Rendón-Salinas, 2014). Destruction of their host plants via herbicide use and reduction of overwintering sites through logging and other disturbances are major contributors to the decline (Flockhart et al., 2013; Pleasants & Oberhauser, 2013).

Monarchs are dependent upon milkweed plants of the family Apocynacea, primarily the genus *Asclepias*, for breeding. More than 130 milkweed species are found in North America (Luna & Dumroese, 2013), although their distribution, abundance, and suitability vary considerably (Lemoine, 2015). The butterfly’s tight association with milkweeds reflects their dependence on plant cardenolides which they sequester for defense (Reichstein, Euw, Parsons, & Rothschild, 1968) and their ability to overcome the obstacles presented by the latex emitted by the plants in response to larval feeding (Agrawal & Konno, 2009). Milkweed species vary not only in their relative levels of cardenolides and latex but also in their relative attractiveness for oviposition by female monarchs (Ladner & Altizer, 2005). In addition, larval growth and development differs with host plant species (Erickson, 1973). Monarchs encounter different milkweed species along their migratory route because phenology and geographical distributions vary between species of milkweeds (Dingle, Zalucki, Rochester, & Armijo-Prewitt, 2005). Females thus may encounter situations in which the best host species is not available, forcing them to oviposit on the milkweed species present. On the other hand, a non-migratory population may become adapted to the most prevalent host plant in its area.

*Asclepias curassavica,* or tropical milkweed, originated in Mexico (Woodson, 1954) but has become invasive in many parts of the world (Ward & Johnson, 2012). This species has been implicated, because it provides a breeding site throughout the year, in discouraging monarchs from migrating, leaving them vulnerable to infection by parasites (Satterfield, Maerz, & Altizer, 2015). Also invasive is *Gomphocarpus physocarpus* (formerly *Asclepias physocarpa*), native to Southeast Africa (Notten, 2010), and utilized in Hawaii and Australia where non-migratory monarch populations have become established. *Asclepias linaria* is native to North America, found in most states of Mexico and in the Southwestern United States (Global Biodiversity Information Facility http://www.gbif.org/species/3170270). The loss of monarch habitat at the same time that invasive milkweeds are providing novel host plants, raises questions about the butterflies’ ability to utilize this new range of plants, and, if so, what genes underlie the ability to switch hosts.

Near Irapuato, in the Mexican state of Guanajuato, a nonmigratory population of monarchs is associated with *A. curassavica* (Pfeiler et al., 2017). Although the length of time that this population has existed is unknown, it shows genetic differentiation from the migratory populations in MBBR (Pfeiler et al., 2017) and thus may have become adapted to its local *A. curassavica* hosts. Depending upon the extent of adaptation, and the chemical differences characterizing other milkweeds, the Irapuato population could exhibit fitness decrements on alternative hosts. We thus explored performance, together with transcriptional differences, of monarch larvae from Guanajuato reared on three different milkweed hosts. We reared second instar larvae for 24 hours on leaves of *A. curassavica* as well as the other two milkweeds mentioned above: *A. curassavica and G. physocarpus.* We measured their sizes before and after feeding and examined their transcriptomic responses, both in coding genes and small noncoding RNAs, to feeding upon these three hosts. We expected their growth would be highest on *A. curassavica,* followed by *A. linaria* with the poorest performance on *G. physocarpus* a species not yet reported in Mexico. Furthermore, we expected larger transcriptional differences between *G. physocarpus* and any of the *Asclepias* than between *Asclepias* given the correlation between phylogeny and chemical diversity (Agrawal, Salminen, & Fishbein, 2009; Rønsted et al., 2012). Concomitant with the effects on growth, we expected the greatest disturbance in gene expression to be observed for larvae having fed on *G. physocarpus,* relative to the other two plants.

## 2. MATERIALS AND METHODS

### 2.1 Collection and handling of monarch eggs and larvae

Monarch eggs were collected from leaves of *A. curassavica* wild plants and kept in the laboratory on the same leaves, in petri dishes, until larvae emerged. Once the larvae emerged we transferred them to fresh *A. curassavica* leaves until they achieved the 2^nd^ instar stage. We then transferred early 2^nd^ instar larvae to fresh leaves of the three different *Asclepias* species (*A. curassavica, A. linaria* and *G. physocarpus*) and allowed them to feed for 24 hrs. A subset was utilized to measure larval size before and after feeding (growth) and another subset was used for the differential gene expression study.

### 2.2 Host plants

Plants of *A. curassavica, A. linaria,* and *G. physocarpus* were reared in the greenhouse at Langebio under temperature and relative humidity ranging diurnally between 25-28C and 42-78%, respectively. Mature leaves were placed in petri dishes lined with moist towels and second instar larvae were placed on the leaves to feed for 24 hours in the laboratory at 24C, and 12hL:12hD.

### 2.3 Measuring development

The length, width and head size of individual larvae were measured before they were placed on the leaves to feed and again after 24 hours. Head size was used to confirm developmental stage (between 0.8mm and 1.5mm for second instar). Larval growth during the 24 hours was the difference in size, in cubic millimeters, before and after feeding on the different host plants.

### 2.4 RNA extraction and sequencing

Larvae were collected after 24 hours, frozen immediately in liquid nitrogen and stored at −80C until the total RNA extractions were performed. Triplicates of five whole larvae for each treatment were homogenized using a Tissue Lyzer. Total RNA was extracted using the Direct-zol kit (Zymo), using the same total RNA to prepare mRNA and sRNA libraries. The mRNA libraries were created using TruSeq RNA kit (LANGEBIO, CORE-FACILITY) and sequenced on an Illumina HiSeq 2500 platform using 2×150 paired-end format (GENEWIZ). The small RNA libraries were created using the Illumina TruSeq small RNA kit and sequenced on an Illumina HiSeq 2500 with a 1×50 single-end format (GENEWIZ).

For each diet we prepared three libraries, each being the pool of 5 larvae fed with the same diet, which we consider as biological replicates. Individual mRNA libraries yielded between 12.8 and 16.9 million paired-end reads, for a total of 131.7 million (Supp. Table 1). The small RNA libraries yielded between 24.5 and 35.3 million reads, for a total of 275.9 million (Supp. Table 2).

### 2.5 Analysis of RNA-Seq experiments

Adapter and low quality bases (mean PHRED score < 15 in windows of 4 nucleotides) were removed with Trimmommatic (Bolger, Lohse, & Usadel, 2014). Cleaned reads shorter than 30 nucleotides long were discarded. Filtered reads were aligned against the monarch butterfly genome (MonarchBase, v3 assembly; Zhan and Reppert, 2013) using HISAT2 v2.1 (maximum multimapping sites = 900, --max-intronlen 15000; Pertea, Kim, Pertea, Leek, & Salzberg, 2016). Gene counts were summarized with FeatureCounts (Liao, Smyth, & Shi, 2014) using the official gene set v2 (OGS2) from MonarchBase. Paired reads were considered as a single count. After quality filtering and adapter trimming, 92%-96.4% of the paired-end reads were successfully mapped to the genome, and out of these, 82.7-87% were unambiguously assigned to annotated protein-coding genes (Supp. Table 1). Out of the 15,128 annotated genes in the monarch genome, 11,065 were robustly detected as expressed in our experiment (CPM > 1 in at least three libraries, Supp. Table 3).

Differential expression was calculated with edgeR v3.18.1 (Robinson, McCarthy, & Smyth, 2009). Genes with zero counts per million (CPM) in more than five libraries were ignored in downstream analyses. Read counts were normalized by the trimmed mean of M-values (TMM) method. Mean-variance relationship was modeled using voom with quality weights (Liu et al., 2015) through limma v3.32.2. Gene Ontology (GO) and KEGG pathway enrichments were calculated with topGO and kegga, respectively, using the functional annotation from MonarchBase and UniProtKB. Enriched GO categories were filtered by redundancy with REVIGO (simRel >= 0.5; Supek, Bošnjak, Škunca, & Šmuc, 2011).

### 2.6 Annotation of untranslated regions for protein coding genes

Read alignments by HISAT2 were used for genome guided transcriptome assembly with StringTie (Pertea et al., 2015) disregarding OGS2 annotation. Transcripts overlapping more than one gene were discarded to avoid gene fusions. Filtered transcripts were used in PASA2 pipeline (Haas et al., 2003) for two rounds of gene model updates of OGS2.

### 2.7 Annotation of non-coding RNAs

To identify tRNAs we used tRNAscan-SE v1.23 (Lowe and Eddy, 1996) with default parameters. For identification of 5.8S, 18S and 28S ribosomal RNAs we used RNAmmer v1.2 (Lagesen et al., 2007). For a wider variety of non-coding RNAs, including rRNAs, tRNAs and miRNAs, we used cmsearch from infernal v1.1.1 (Nawrocki & Eddy, 2013) with default parameters and the covariance models from Rfam v12.0 (Nawrocki et al., 2015).

### 2.8 miRNA annotation

Following the approach used in miRBase (Kozomara & Griffiths-Jones, 2014), we defined a cluster as miRNAs on the same strand and separated by less than 10kb. Using Mapmi v159-b32 (Guerra-Assunção & Enright, 2010), we looked for miRBase high confidence mature miRNAs as well as miRNAs identified in *Cameraria ohridella* and *Pararge aegeria* (Quah, Breuker, & Holland, 2015). For identification of precursors of conserved miRNAs and *de novo* miRNA predictions, we relied on miRDeep v2.0.0.8 (Friedländer et al., 2012). Novel predictions were included as novel miRNAs only if they had miRDeep2 scores >100 and were consistently detected in at least 3 libraries.

### 2.9 miRNA target prediction

The updated 3’ UTRs that we produced in this work were used to predict targets for both potential mature miRNAs from each miRNA loci. Target prediction was performed with TargetScan 7 (Agarwal, Bell, Nam, & Bartel, 2015) without conservation information and requiring a context++ score < −0.6.

### 2.10 Analysis of sRNA-seq experiments

Using reaper (Davis, van Dongen, Abreu-Goodger, Bartonicek, & Enright, 2013), we trimmed the raw short reads at the start of the 3’ adapter (TGGAATTCTCGGGTGCCAAG), or if the median Phred score of four nucleotides went below 15 or if the dust complexity score went below 20 (e.g. to remove runs of a single nucleotide at the end of a read). After trimming, reads shorter than 11 bases were discarded. The length distribution of the sequenced small RNA reads exhibited a clear peak around 22-23 for the *G. physocarpus* libraries and one replicate of *A. linaria* as would be expected for mature miRNA sequences. Although noisier, the two other replicates of *A. linaria* and one of *A. curassavica* also showed the same trend. Two libraries from larvae fed on *A. curassavica* had a more homogenous distribution, suggesting more extensive RNA degradation (Supp. Figure 1).

We aligned the trimmed reads with ShortStack v3.8.2 (Axtell, 2013) using align_only and nostitch options. Briefly, ShortStack uses bowtie (Langmead, Trapnell, Pop, & Salzberg, 2009) for read alignment allowing zero mismatches and a maximum of 500 multiple mapping sites per read. On average, 61.9% of the filtered reads aligned to the genome, varying between 87% and 27% (Supp. Table 2). Most of the aligned reads not mapping to miRNAs were assigned to rRNAs or mRNAs, consistent with RNA degradation being the main reason for the differences in the distributions.

We summarized the mature miRNA counts with featureCounts requiring reads to map on the same strand as the miRNA. To test for differences in gene expression, we used edgeR v3.18.1 (Robinson et al., 2009). Genes with zero counts per million (CPM) in more than five libraries were ignored in downstream analyses. Read counts were normalized by the trimmed mean of M-values (TMM) method.

## 3. RESULTS

### 3.1 Larval growth

Larval growth was calculated as the volume difference before and after 24 hours of feeding on the different hosts (Figure 1). All larvae survived on all three milkweed species but grew at different rates depending on host (ANOVA: df = 2, F = 14.96, P < 0.0001, Supp. Table 5). Larvae grew the most on *A. linaria* followed by *A. curassavica*, although the difference between these two, after transforming the data, was not significant. Larvae on *G. physocarpus*, however, barely grew at all (Supp. Table 6).

**Figure 1.**
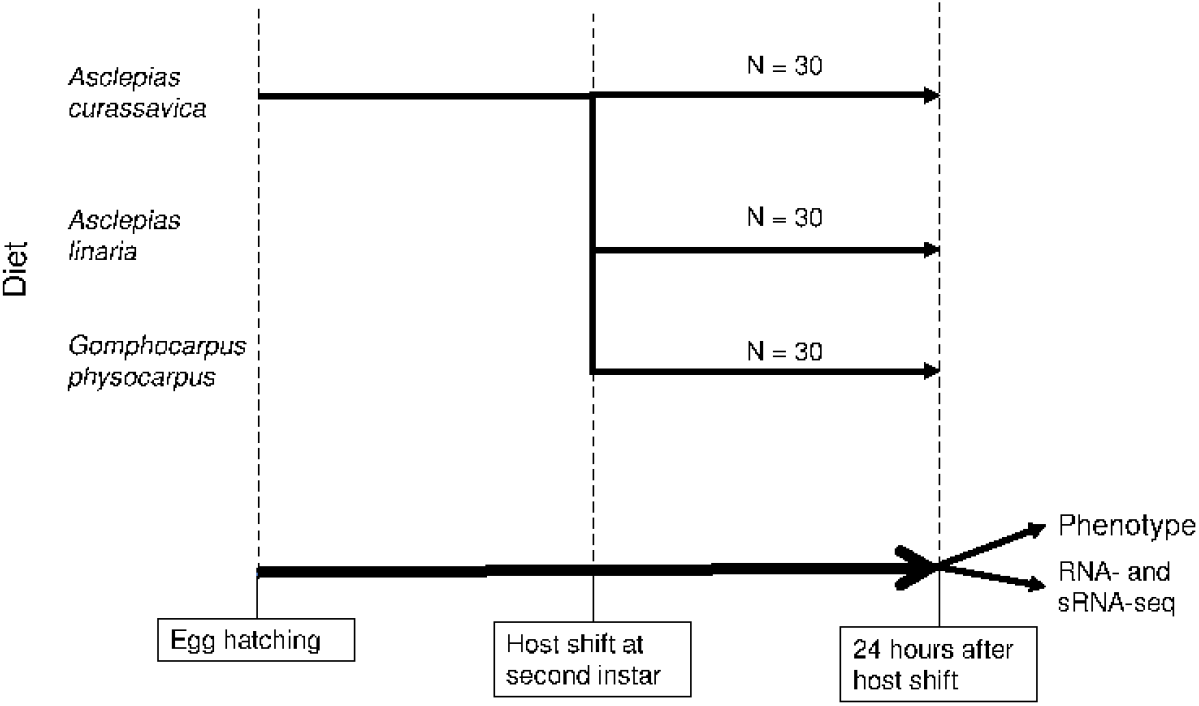
*Danaus plexippus* larval host shift. Larvae collected and reared on *A. curassavica,* after reaching the second instar, were shifted to *A. linaria* or *G. physocarpus* or remained feeding on *A. curassavica*. Volume was measured at host shift and 24 hours later, while RNA was extracted at the end of the treatment.

### 3.2 Differential expression analysis of coding genes

Gene expression profiles of larvae fed on *A. linaria* and *A. curassavica* were quite similar, as only 66 protein-coding genes were differentially expressed (DE) between them and all of them had higher abundance in larvae fed on *A. linaria* (Supp. Table 7). When these two *Asclepias* are compared to *G. physocarpus*, however, we found 5,811 DE genes (FDR < 0.05) for larvae fed on *A. curassavica*, and 4,522 for those fed on *A. linaria* (Figure 2A; Supp. Table 8). Of these two comparisons, there were 3,896 DE genes in common (Figure 2B). The differential expression results correlate with larval growth differences, in both cases there was little difference between larvae fed on *A. linaria* and those fed on *A. curassavica*, while those fed on *G. physocarpus* were the most different. Considering this, we decided to focus on the expression profile of larvae fed on *G. physocarpus*, compared to the *Asclepias* diets.

**Figure 2.**
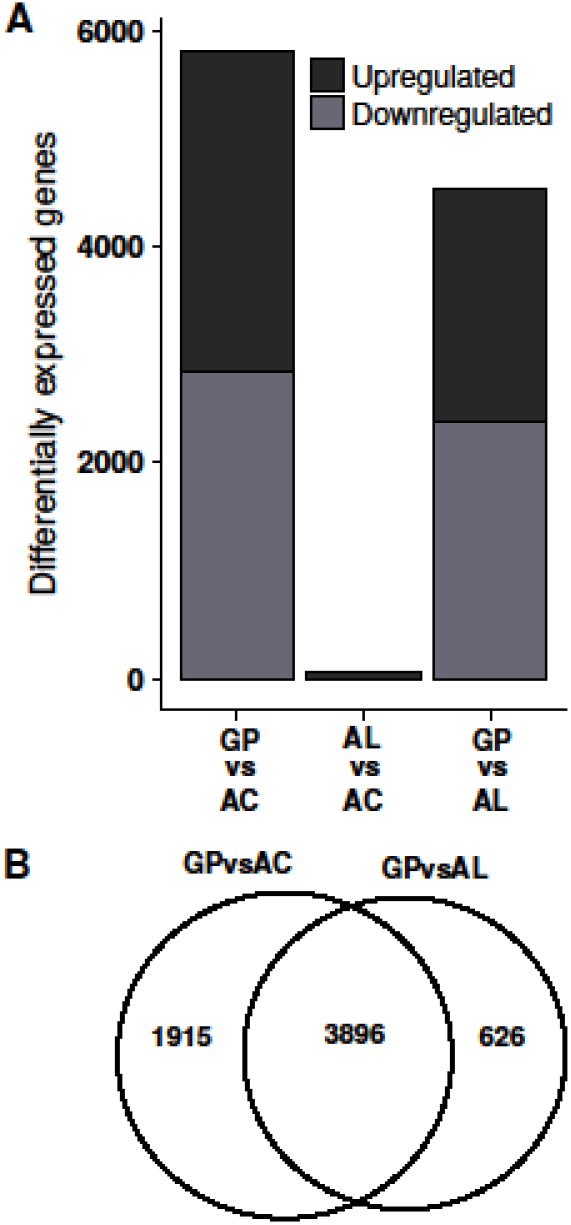
Differentially expressed protein coding genes. A) Number of differentially expressed genes identified in each diet comparison. B) Genes found differentially expressed in both or one contrast using larvae fed on *G. physocarpus* as the reference. FDR < 0.05.

### 3.2.1 Functional enrichment analysis

The *G. physocarpus* vs *A. curassavica* comparison showed many of the same pathways enriched in the *G. physocarpus* vs *A. linaria* contrast (Table 1), as expected due to the high overlap of differentially expressed genes. The most affected pathways were related to growth and detoxification. In larvae fed on *G. physocarpus,* cell cycle and proteasome KEGG pathways were down-regulated while ATP binding cassette (ABC) transporters were up-regulated. Analyses using Gene Ontology (GO) revealed similar results, including down-regulation of terms related to DNA repair and replication, cell cycle and the proteasome, while cation transport and G-protein signaling functions were up-regulated compared to larvae fed on the *Asclepias* (Supp. Table 9).

**Table 1.** Over-represented KEGG pathways among differentially expressed genes in both contrasts taking as reference larvae fed on *Gomphocarpus physocarpus*.

| Pathway | Direction | N | Up CP | Up LP | Down CP | Down LP | FDR CP | FDR LP |
| --- | --- | --- | --- | --- | --- | --- | --- | --- |
| DNA replication | Down | 41 | 1 | 1 | 38 | 34 | $5.29 \times 10^{-16}$ | $6.65 \times 10^{-12}$ |
| Protein processing in endoplasmic reticulum | Down | 154 | 18 | 12 | 88 | 84 | $1.93 \times 10^{-12}$ | $4.37 \times 10^{-12}$ |
| Proteasome | Down | 46 | 1 | 1 | 37 | 38 | $3.12 \times 10^{-11}$ | $2.47 \times 10^{-13}$ |
| Spliceosome | Down | 162 | 22 | 17 | 87 | 78 | $3.48 \times 10^{-10}$ | $1.44 \times 10^{-7}$ |
| Aminoacyl-tRNA biosynthesis | Down | 53 | 3 | 2 | 36 | 32 | $2.81 \times 10^{-7}$ | $2.66 \times 10^{-5}$ |
| Cell cycle | Down | 99 | 12 | 10 | 56 | 54 | $2.92 \times 10^{-7}$ | $2.39 \times 10^{-7}$ |
| Nucleotide excision repair | Down | 49 | 3 | 1 | 32 | 25 | $1.01 \times 10^{-5}$ | $3.1 \times 10^{-2}$ |
| Mismatch repair | Down | 23 | 1 | 0 | 19 | 18 | $1.48 \times 10^{-5}$ | $4.98 \times 10^{-5}$ |
| Protein export | Down | 23 | 0 | 1 | 18 | 17 | $1.50 \times 10^{-4}$ | $4.35 \times 10^{-4}$ |
| Gastric acid secretion | Up | 55 | 29 | 24 | 9 | 7 | $1.19 \times 10^{-2}$ | $2.13 \times 10^{-3}$ |
| Salivary secretion | Up | 71 | 35 | 28 | 12 | 10 | $1.33 \times 10^{-2}$ | $4.06 \times 10^{-3}$ |
| ABC transporters | Up | 53 | 28 | 21 | 3 | 3 | $1.51 \times 10^{-2}$ | $3.96 \times 10^{-2}$ |
| Phototransduction | Up | 30 | 18 | 18 | 5 | 3 | $3.56 \times 10^{-2}$ | $8.18 \times 10^{-5}$ |

Over-represented KEGG pathways (FDR < 0.05) among differentially expressed genes (FDR < 0.05) when contrasting larvae fed on *A. curassavica* vs *G. physocarpus* (CP) or *A. linaria* vs *G. physocarpus* (LP). Column 'N' indicates the total number of assessed genes which are member of that pathway while 'Up' and 'Down' shows the number of significantly up- or down-regulated, respectively, genes in each contrast. The 'FDR' columns present the significance, corrected for multiple testing, of the category for the direction, up or down, with more differentially expressed genes.

#### 3.2.1 Deregulated genes

Several well-studied genes related to growth and the cell cycle were up-regulated in larvae fed on *G. physocarpus,* including the fork head box transcription factor *foxo*, the Insulin-like receptor *InR* and the inhibitor of translation initiation factor eIF4E (4E-BP), *thor*. Consistently, *thor* was downregulated. In agreement with reduced translation, most subunits of the proteasome were downregulated. Down-regulated cell cycle genes included cyclins A, B, D and E. The central kinase of the Hippo signaling pathway, *hpo*, was down-regulated, while *yorkie*, a transcriptional co-activator repressed by Hippo signaling, was up-regulated (Supp. Table 10).

Even though they do not belong to a single pathway, many detoxification related genes were up-regulated in larvae fed on *G. physocarpus*, including 11 alcohol dehydrogenases (ADHs), 15 carboxylesterases (CEs), 12 glutathione S-transferases (GSTs), 33 cytochromes P450 (P450s) and 12 UDP-glucuronosyltransferases (UGTs) (Supp. Table 11). On the other hand, taken as a whole, the superfamily of ATP-binding cassette (ABC) transporters was significantly enriched among up-regulated genes. Out of 55 ABC transporters in the monarch genome, 28 had significantly greater expression in larvae fed on *G. physocarpus* relative to that of the other diets. Conversely, only two had lower expression. *Mdr49* had the second strongest fold change among ABC transporters, just preceded by *ABCA13*, increasing more than four-fold in larvae fed on *G. physocarpus*.

### 3.3 MicroRNAs in D. plexippus larvae

Together with protein coding genes, regulatory elements such as microRNAs (miRNAs) could be involved in the larval response to diet. MiRNAs are single stranded ~22 nt RNAs that cause mRNA destabilization upon binding through partial complementarity to their 3’ UTR. Using complementary expression- and homology-based approaches, we identified 99 miRNAs in the monarch genome, including 87 conserved ones, 3 previously reported as monarch-specific and not previously reported in miRBase, and 9 novel miRNAs (Supp. Table 12). Although conserved miRNAs could be identified independently of gene expression, all the 99 miRNAs were expressed in at least 5 libraries. Furthermore, we identified miR-12 from the miR-12/304/283 cluster which was previously unidentified in the monarch genome.

We next examined the expression levels of mature miRNAs and found that the most highly expressed miRNAs were miR-281 and miR-31 with close to 20% and 14% of total reads, respectively, followed by miR-6094, miR-10, miR-8, miR-263a and bantam (Supp. Figure 2).

#### 3.3.1 Differential expression analysis of miRNAs

Our analyses of miRNAs comparing monarchs fed on *G. physocarpus* vs *A. linaria* only identified miR-278-5p and miR-novel37-5p as differentially expressed, with lower and higher, respectively, abundance in larvae fed on *G. physocarpus*. The contrast between larvae fed *A. curassavica* vs the ones fed *A. linaria* found DE miR-novel36-5p, miR-novel28-3p and miR-6094-3p. Comparing the ones fed on *G. physocarpus* and *A. curassavica* diets identified 49 DE miRNAs, including miR-278-5p (Supp. Table 13). Twenty-six of these DE miRNAs have higher abundance in larvae fed *G. physocarpus*, including both arms of miR-2a-1, miR-277, miR-278, miR-1175 and miR-6094. Finally, most of the DE miRNAs are the predominantly expressed arm, except for bantam, miR-9a, miR-9c, miR-10, miR-92b, miR-275, miR-276, miR-279b, miR-306 and miR-965.

#### 3.3.2 Predicted targets for deregulated miRNAs

Since miRNAs are negative regulators of protein-coding genes, we need to identify their targets to understand their functions. Of all potential targets, we focused on coherent targets: those that changed in the opposite direction to the miRNA (Supp. Table 14). For example, we found a complementary site for miR-278-3p in the 3’ UTR of expanded (*ex*), and we observed that this gene was up-regulated while miR-278 was down-regulated in larvae fed on *G. physocarpus*. Similarly, *kibra* and Death-associated inhibitor of apoptosis 1 (*diap1*) are coherent targets of miR-278-3p.

Finally, we assessed enrichment of KEGG pathways to predict the functional categories that could be affected by differential miRNA regulation (Supp. Table 15). Hippo signaling was enriched among the coherent targets of miR-2755-5p, miR-278-5p and miR-278-3p. ABC transporters were enriched among miR-2a-1-3p, miR-2a-1-5p, miR-92b and miR-1175-5p. In addition, insulin signaling was enriched among miR-100 and miR-1175-3p targets. miRNAs whose targets were enriched in cell cycle are miR-novel42, miR-10, miR-14, miR-745, miR-927a, miR-970 and miR-993.

## 4. DISCUSSION

### 4.1 Performance differences

While we predicted relatively poor performance on *G. physocarpus* relative to the two *Asclepias*, the near lack of growth was surprising. Several factors could have influenced this outcome. One is that early feeding experience influenced subsequent consumption (Hu et al., 2018; Jermy, Hanson, & Dethier, 1968). We had kept first instar larvae on the *A. curassavica* on which the eggs were laid for 24 hours until they reached second instar, in order to avoid damaging the fragile eggs or first instar larvae. Thus, second instar individuals were either retained on *A. curassavica* for another 24h as second instars or, transferred to either *A. linaria* or *G. physocarpus*. It is not clear if the initial exposure to one diet influenced their feeding and growth if transferred to something different. Clearly this was not the case for the switch to *A. linaria,* which is more closely related to *A. curassavica*, than to *G. physocarpus*. Furthermore, *A. curassavica* and *G. physocarpus* have similar leaf shapes, both lanceolate, while *A. linaria* has them in filiform shape. And yet larvae barely grew on *G. physocarpus* while those on the two *Asclepias* increased their volume by a factor greater than two. Therefore, leaf shape is unlikely to explain the growth differences.

Differences in host quality and chemistry, however, could have slowed development on *G. physocarpus*. In *Danaus plexippus*, larvae consume more host mass when the host is of poorer quality (Lavoie & Oberhauser, 2004). We did not measure consumption, so its role in our results is unclear. Alternatively, *Gomphocarpus physocarpus*, may contain chemicals toxic to the young larvae, as suggested by the differential expression of detoxification gene, as discussed below.

A wide range of differences in the literature, with respect to experimental components and design also are likely to influence the outcomes of various larval feeding experiments. It’s unclear whether allowing the larvae to grow to adulthood or to pupation would have produced a different outcome. Our interest was in short-term exposures of young larvae to the different plants. In previous studies, however, where larvae grew for longer periods, they grew less on *A. linaria* than in *A. curassavica* (Petschenka & Agrawal, 2015; Tao, Hoang, Hunter, de Roode, & Cotter, 2016) or had higher relative survival on *G. physocarpus* (Tao et al., 2016). In these earlier experiments, larvae fed on living plant material, versus the cut leaves we utilized, possibly altering their exposure to the defensive latex compound present in the plant (Agrawal & Konno, 2009). Finally, the population of nonmigratory monarch butterflies from Irapuato Mexico that we used here exhibit genetic differences (Pfeiler et al., 2017) compared to the migratory populations from which the strains in the previous experiments were derived. Experimental design, including source populations of monarchs used and timing of exposure, can influence the effect of host plant on performance. As most monarch feeding experiments were performed differently from one another, direct comparisons are not possible. Thus, while our results only reflect performance for a short period in early larval development, growth in the early instars is likely critical to longer term fitness.

### 4.2 Differential expression of coding genes

Cell cycle, DNA replication and proteasome were enriched among downregulated genes, suggesting a molecular orchestration of growth arrest for *G. physocarpus*. In agreement with the performance results, larvae fed on *G. physocarpus* had lower expression of cell cycle and growth regulation genes from the insulin pathway. As discussed above, arrested growth could have been caused by lack of palatability, digestibility, nutrition, or interference by some unknown plant compound. Although both transcriptomic and performance results are consistent with lower nutritional intake, we found no enrichment for carbohydrate or lipid metabolism genes. This is hard to explain given that, in *Bombyx mori*, 3 hours of starvation are enough to induce the hyperglycemic response (Satake, Kawabe, & Mizoguchi, 2000). If the G*. physocarpus* larvae were starving, we would expect that at 24 hours lipid metabolism would be more active relative to the other diets and that carbohydrate metabolism genes would be less active. But this was not the case. Only when comparing *A. linaria* vs *A. curassavica* was there a slight up-regulation of genes involved with carbohydrate, amino acid and fatty acid metabolism in larvae fed on *A. linaria*.

Other grow-related processes were more consistently affected. Inhibition of the proteasome correlates with growth arrest (Yin et al., 2005), and the components of the proteasome were down-regulated in larvae fed on *G. physocarpus* compared to those fed on the *Asclepias*. *Tor* is a well-known regulator of growth size in response to dietary intake. Tor requires raptor to regulate eukaryotic translation initiation factor 4E-binding protein (Thor) by phosphorylation (Hara et al., 2002). Although *Tor* itself was not differentially expressed, regulatory-associated protein of Tor (*raptor*) was significantly upregulated. Another important pathway for growth regulation is Hippo signaling which restrains cell proliferation (Harvey, Pfleger, & Hariharan, 2003). Inactivation of Hippo kinase results in translocation of yorkie to the nucleus where it transcribes genes involved in cell proliferation in mammals and *Drosophila* (Meng, Moroishi, & Guan, 2016). Consistently, *hippo* was down-regulated and *yorkie* up-regulated in larvae fed on *G. physocarpus*. As part of the insulin signaling pathway, transcription factor forkhead box sub-group O (*foxo*), up-regulated in larvae fed *G. physocarpus*, regulates the effect of nutrition on *Drosophila* organ size (Tang, Smith-Caldas, Driscoll, Salhadar, & Shingleton, 2011). Serine/threonine protein kinase Akt phosphorylates foxo leading to its cytoplasmic retention rendering it unable to regulate its targets. When stressed by starvation, foxo relocalizes to the nucleus and activates two key players of the Akt/PKB signaling pathway: *Thor* and insulin receptor (*InR*) (Puig, Marr, Ruhf, & Tjian, 2003). *Thor* was strongly up-regulated while *InR* was slightly down-regulated in the *G. physocarpus* fed larvae. Our results are consistent with these larvae being under nutritional stress and not growing.

Growth could also be impaired by toxicity of unknown plant compounds. Detoxification has been subdivided into three phases (Brattsten, 1988). The first one directs catabolism of xenobiotics, while the second phase conjugates those products, and in the final phase these more hydrophilic compounds are excreted or sequestered. While genome-wide expression studies of host shifts in other insect taxa revealed differential expression of detoxification genes involved in detox phases I and II (De Panis et al., 2016; Govind et al., 2010; Ragland et al., 2015), we found no enrichment of these categories in any contrast. Detoxification genes involved in phase lll, namely the ABC transporters, however, are increasingly considered relevant for host shifts (Bretschneider, Heckel, & Vogel, 2016; Koenig et al., 2015) and can participate in allelochemical sequestration (Strauss et al., 2014; Strauss, Peters, Boland, & Burse, 2013). Most ABC transporters were significantly overexpressed while only two reduced their expression in larvae fed *G. physocarpus* relative to that of the *Asclepias*. This raises the question of what plant defenses would elicit only transmembrane transport without requiring an increase of its solubility or conjugation by the herbivore to cope with it. Among the well-known milkweed defenses are toxic cardiac glycosides to which *Danaus plexippus* is immune, but modulates the concentration it sequesters (Holzinger, Frick, & Wink, 1992; Petschenka & Agrawal, 2015; Vaughan & Jungreis, 1977). One of these upregulated ABC transporters, multidrug resistance protein 49 (*Mdr49*), has been found to alleviate the toxic effects of cardenolides in *D. melanogaster* (Groen et al. 2017) suggesting that some of the ABC transporters upregulated might be linked to increased cardenolide transport.

### 4.3 Differential expression of microRNAs and their targets

Since miRNAs can detectably repress their targets at the transcript level, we followed the convention of selecting only the predicted targets that change in the opposite direction to the miRNA. Nevertheless, miRNA profiling in whole organisms can be problematic because many miRNAs are only expressed during particular developmental stages and/or in specific cell types. Obviously, their repressive function can only be exerted in the cells in which they are expressed. Predicting miRNA targets amongst transcripts expressed in whole organism samples thus may yield many false predictions if the miRNA/target pair are not expressed in the same cells. Conversely, if the miRNA is expressed in just a small subset of the sampled tissue, while the target is more generally expressed, the effect on its targets can easily be undetectable.

Unexpectedly, we found very few differentially expressed miRNAs when comparing larvae fed on *G. physocarpus* and *A. linaria*, although our coding gene analysis showed a drastic difference between these two diets. This is likely a consequence of the higher variation between the *A. linaria* miRNA libraries (Supp. Figure 3). Although the same RNA was used for both sequencing strategies, the mRNA libraries from larvae fed on *A. linaria* showed less variability. Hence, due to reduced statistical power, there might still be several unidentified differentially expressed miRNAs in this comparison. Despite this, we found miR-278 to be consistently down-regulated in the larvae fed on *G. physocarpus,* making it the most obvious candidate for follow-up analyses. This miRNA might be relevant in the monarch-milkweed interaction because, in *D. melanogaster*, null mutants of this miRNA are defective in energy homeostasis, leading to a lean phenotype (Teleman & Cohen, 2006) while its overexpression results in eye and wing overgrowth (Nairz et al., 2006). Furthermore, we predicted three genes of the Hippo signaling pathway: *ex*, *kibra* and *diap1* to be up-regulated targets of this miRNA. Kibra and ex, together with Merlin (Mer), form a complex acting upstream of yorkie, which in turn regulates *Diap1* (Genevet, Wehr, Brain, Thompson, & Tapon, 2010).

Many studies of microRNA targets focus only on the annotated mature miRNA (produced from one of the hairpin arms). However, here we found that some miRNAs show high expression of both 5p and 3p arms, both of which can be differentially expressed. This is important, since the two arms of a miRNA generally have different “seed” sequences, and they rarely target overlapping sets of genes (Marco, MacPherson, Ronshaugen, & Griffiths-Jones, 2012). In the case of miR-278, both arms are down-regulated, and each arm targets different genes of the Hippo pathway Thus, miR-278, when active, could be one of the main upstream regulators of a growth response. In our experiments, larvae fed on *G. physocarpus* have lower levels of miR-278, consistent with the general signatures of growth inhibition.

Even though our results regarding miRNA differential expression already suggest new growth regulating interactions, sampling of more homogenous tissues or cells would be useful to gain cleaner measurements of miRNAs and their targets. Further experimental validation will also be required to demonstrate a direct effect of each miRNA and its predicted targets, in the context of the function we describe here.

## 5 CONCLUSION

In conclusion, a growth period of 24 hours for second instar on three different milkweed hosts produced significant differences in performance and expression patterns in coding and noncoding RNAs. Both performance and expression were the most similar on the most closely related milkweed hosts. The monarch larvae we utilized came from a local non-migratory population in Irapuato, Guanajuato, Mexico where *G. physocarpus* has yet to be introduced and where the predominant milkweed appears to be *A. curassavica*. As *G. physocarpus* is reported to be utilized successfully by monarchs in Australia, the differences we observe could reflect a degree of local adaptation as well as the fact that the two *Asclepias* are more chemically similar to each other than to *G. physocarpus*. Monarch host plants are changing in their global distributions, with some favored species disappearing and others invading the migratory routes. The survival of the monarchs depends, in part, upon their ability to successfully reproduce on new host plants. This ability, in turn, depends upon the existence of genetic variability at the loci underlying host use and shifts. Our study identified a number of differentially expressed genes and microRNAs that may influence the future success of the monarchs in the face of global change. An important next step is to determine the diversity in these genes in monarchs from different parts of their range and evaluate this variability or its absence in the context of conservation efforts.

## Supporting information

List of Supplemental Tables

## ACKNOWLEDGEMENTS

This work was supported by a grant from the Alianza Carlos Slim to TAM, a CONACyT graduate fellowship to PMGdlR (598430), funds from Langebio-Cinvestav to CA-G and TAM.

## DATA ACCESSIBILITY

PolyA-mRNA and small RNA Illumina reads and our microRNA annotation file can be found in the Gene Expression Omnibus (GEO) under the accession GSE120501.

## AUTHOR CONTRIBUTIONS

MRLL, TAM and CAG designed the experiments; MRLL performed the growth experiments and extracted RNA for sequencing; PMGdlR and TAM analyzed the growth results; PMGdlR performed all the bioinformatic analyses; PMGdlR, TAM, and CAG interpreted the results and wrote the manuscript.

