## Supplemental Figures for "Differential gene expression reflects larval development and survival of monarch butterflies on different milkweed hosts"

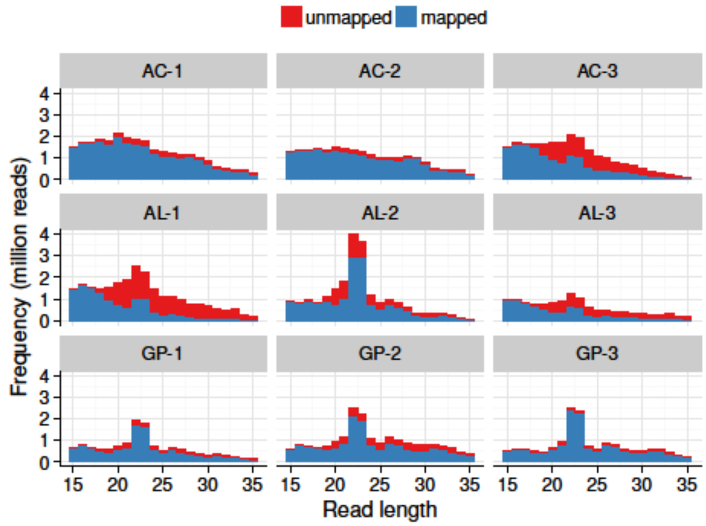


**Supp. Figure 1.** Mapping rate by read length. One mismatch was allowed.

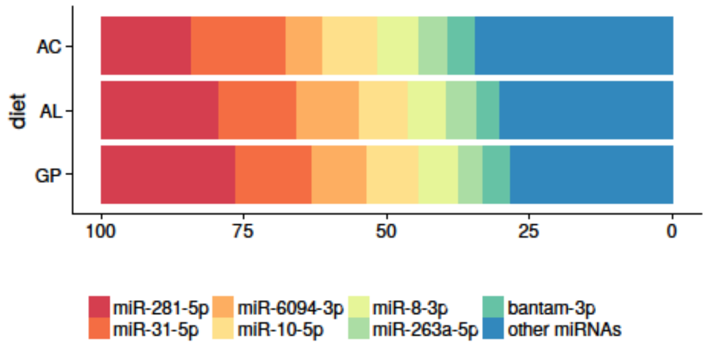


**Supp. Figure 2.** Top seven expressed mature miRNAs. The average fraction of the three biological replicates is shown for each diet.

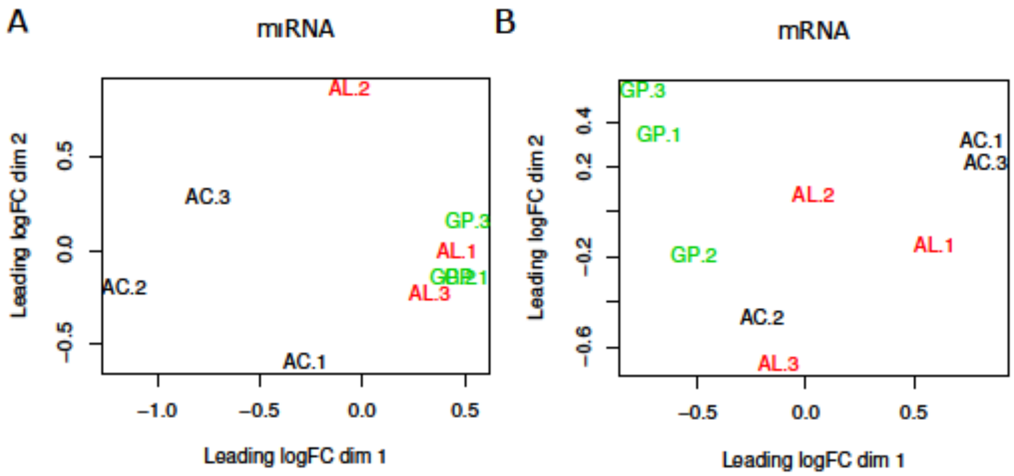


**Supp. Figure 4.** Multi-dimensional Scaling plots showing relationships for sRNA-Seq samples (A) or mRNA-Seq samples (B). Larva were fed on *A. curassavica* (AC), *A. linaria* (AL), or *G. physocarpus* (GP). Numbers indicate biological replicates.
