## Supplementary material for "Differential gene expression reflects larval development and survival of monarch butterflies on different milkweed hosts": List of Supplemental Tables

### SUPPLEMENTARY TABLES

**Supp. Table 1.** RNA-seq summary statistics.

**Supp. Table 2.** sRNA-seq summary statistics.

**Supp. Table 3.** Counts per million of protein coding genes.

**Supp. Table 4.** Genes differentially expressed between larvae fed on *Asclepias linaria* vs *Asclepias curassavica*.

**Supp. Table 5.** ANOVA test of differences between square root transformed larval growth ($mm^3$).

**Supp. Table 6.** Tukey's HSD test square root transformed larval growth at 95% family-wise confidence level.

**Supp. Table 7.** Genes differentially expressed in larvae fed on *Asclepias curassavica* relative to *Asclepias linaria*.

**Supp. Table 8.** Genes differentially expressed either between larvae fed on *Asclepias linaria* or *Asclepias curassavica* relative to *Gomphocarpus physocarpus*.

**Supp. Table 9.** Biological processes enriched among genes differentially expressed between larvae fed either on *Asclepias linaria* or *Asclepias curassavica* vs *Gomphocarpus physocarpus*.

**Supp. Table 10.** *Danaus plexippus* ortholog genes of *Drosophila melanogaster* for hippo, foxo, tor, apoptosis and proteasome pathway that were differentially expressed in any contrast involving larvae fed on *G. physocarpus*.

**Supp. Table 11.** Detoxification genes differentially expressed between larvae fed either on *Asclepias linaria* or *Asclepias curassavica* vs *Gomphocarpus physocarpus*.

**Supp. Table 12.** Coordinates, clustering and arm preference of the monarch miRNAs.

**Supp. Table 13.** miRNAs differentially expressed identified in each of the three contrasts. FDR < 0.05.

**Supp. Table 14.** Differentially expressed coherent targets of the differentially expressed miRNAs.

**Supp. Table 15.** KEGG pathways and GO biological processes enriched among coherent targets.
